# Maternally derived sex steroid hormones impact sex ratios of loggerhead sea turtles

**DOI:** 10.1101/2020.01.10.901520

**Authors:** Emma C. Lockley, Thomas Reischig, Christophe Eizaguirre

## Abstract

Global warming could drive species with temperature-dependent sex determination to extinction by persistently skewing offspring sex ratios. Evolved mechanisms that buffer these biases are therefore paramount for their persistence. Here, we tested whether maternally-derived sex steroid hormones affect the sex-determination cascade and provide a physiological mechanism to buffer sex ratio bias in the endangered loggerhead sea turtle (*Caretta caretta*). We quantified estradiol and testosterone in nesting females and their egg yolks at oviposition, before incubating nests in situ at standardised temperatures. Upon hatchling emergence, we developed a new, non-lethal method to establish the sex of individuals. Despite standardised incubation temperatures, sex ratios varied widely among nests, correlating non-linearly with the estradiol:testosterone ratio in egg yolks. Males were produced at an equal ratio, with females produced either side of this optimum. This result provides evidence that maternal hormone transfer forms a physiological mechanism that impacts sex determination in this endangered species.

## Introduction

Fifty years after the discovery of environmental sex determination, our understanding of its evolutionary significance, underlying mechanisms and ecological consequences in the light of environmental change remains incomplete^1–5^. Most reptile and some fish species undergo temperature-dependent sex determination (TSD), in which gonad differentiation is regulated by temperature at a critical period of embryogenesis^6,7^. Some species produce males at moderate temperatures and females at hot and cold extremes (e.g. the American alligator *Alligator mississippiensis*,^8^ Type II TSD), but, more commonly, TSD species produce an increasing proportion of a specific sex across a range of incubation temperatures (Type Ia: Males at low temperatures, e.g. the painted turtle *Chrysemys picta*^9^; Type Ib: Females at low temperatures, e.g. the tuatara *Sphenodon punctatus*^10^). In all cases, both sexes are produced across a transitional range of temperatures, centered on a pivotal temperature at which both sexes develop in equal proportions. While the pivotal temperature varies among clutches within a population^11^, studies have generally focused on quantifying population-level means as important proxies to estimate sex ratios and predict population dynamics^4,12,13^. These population-level proxies suggest that rising global temperatures present the potential for extreme sex ratio biases in TSD species, with implications for population dynamics and extinction risk^4^.

The adaptive value of TSD is still debated, but fitness advantages under sex-specific thermal environments are predicted by the more favoured Charnov-Bull theory of differential fitness^14,15^. The postulates of this theory have been demonstrated in eggs of the Jacky dragon (*Amphibolurus muricatus*) that were experimentally treated with an aromatase inhibitor, constraining embryos to develop as males at female producing temperatures. These males showed lower lifetime reproductive success than controls^3^. While demonstrating the adaptive value of TSD, the use of an aromatase inhibitor to manipulate sex in this study also highlights the role of sex steroid hormones on the TSD mechanism^16–18^.

Exogenous application of estradiol (E_2_) to the incubating eggs of some species can feminise TSD embryos incubated at male-producing temperatures^19–22^. In addition, the application of testosterone (T), the precursor androgen of E_2_, can also feminise embryos via the synthesis of E_2_ from T by the aromatase enzyme ^20,23^, and indeed the use of aromatase inhibitors can force male development^24,25^. Both temperature and exogenous treatment with E_2_ activate the same molecular pathways, altering the transcription of the chromatin modifier gene *Kdm6b,* and conferring sensitivity to a sex-determining gene, *Dmrt1*^2^.

In model TSD species that exhibit Type 1a TSD, such as the slider (*Trachemys scripta*) and the painted turtle, maternal transfer of sex steroid hormones into eggs varies seasonally^16,26^. Elevated concentrations of maternal investment in yolk E_2_ and greater E_2_:T ratios increase the likelihood of feminisation at a given temperature, effectively reducing the pivotal temperature of a clutch by providing substrate that will prime the activation of female-producing molecular pathways^16,26^. Should these patterns be found in non-model species, variation in maternal hormone transfer to eggs could be a universal mechanism to (i) change the threshold at which temperature affects an individual’s sex development, (ii) modify the sex ratio of the clutch, and (iii) possibly buffer against the negative effects of rapid global temperature increase.

There is a particular need to understand the impacts of climate change on the demographics of threatened species. As a consequence of rising temperatures, extreme feminisation of sea turtle populations has been forecast by the end of the century^4,27–29^. Some studies suggest effects are already visible in adult populations^28^. There has been much study into how behavioural responses, such as modified phenology or nest site selection, may mitigate the effects of a warming environment^30,31^, but no overall trends preventing extirpation are visible. There has been, however, little consideration for physiological mechanisms that may increase variation in the pivotal temperature, and therefore on sex ratio. Understanding possible physiological mechanisms has been constrained in sea turtles in particular by the lack of non-lethal methods to sex neonates (but see^32^). This issue is especially important for endangered populations, where sacrificing individuals is not possible.

Here, we tested whether maternally-derived sex steroid hormones affect the sex-determination cascade and the resulting offspring sex ratios in an endangered sea turtle population. Focusing on loggerhead turtles (*Caretta caretta*) nesting in the Cabo Verde archipelago, we standardised the thermal environment of clutches in an experimental field hatchery, exposed to natural conditions. Should temperature be the sole driver of sex determination, similar sex ratios among clutches would be expected under these standardised thermal conditions. Alternatively, any inter-clutch variation would arise from intrinsic characteristics of the eggs, such as maternally-derived hormones. To test these hypotheses, we quantified E_2_ and T concentrations in the plasma of nesting females, their egg yolks and neonates. We developed a non-lethal sexing method using circulatory hormone profiles of neonates, and determined the clutches’ sex ratios. Inter-clutch variation in sex ratio was then linked to yolk hormone concentrations. Finally, we illustrate how maternal hormone transfer impacts sex ratio in the face of IPCC climate change predictions, by re-parameterising a previously used mathematical model^4^ to forecast the future population dynamics of this endangered nesting aggregation.

## Results

This study focused on loggerhead turtles nesting on the island of Boavista in the Cape Verde archipelago. First, using enzyme-linked immunosorbent assays (ELISA – Enzo LifeSciences), we quantified concentrations of the sex steroid hormones E_2_ and T in both the blood plasma of 26 nesting females and up to two of their eggs directly after oviposition. Clutch sizes were recorded at this time. High levels of individual variation were observed in adult plasma hormone levels (SI appendix, Table S1), with a mean T concentration of 1148.48 ± 148.63 (SE) pg/ml, a mean E_2_ concentration of 235.79 ± 22.71 (SE) pg/ml, and a mean E_2_:T ratio of 0.32 ± 0.05 (SE). Linear models (LM) showed positive correlations between E_2_ and T in both the female plasma (SI Appendix, Fig. S1A, F_1,16_ = 4.608, p = 0.048) and their egg yolks (SI Appendix, Fig. S1B, F_1,23_ = 7.338, p = 0.013). In reptiles, maternally derived hormones are constant across all eggs of a given clutch^33^, which we confirmed with a subset of clutches where two egg yolks were analysed (Paired t-tests: T: df = 11, t = 0.224, p = 0.827; E_2_: df = 10, t = −0.885, p = 0.397; E_2_:T: df = 9, t = −1.173, p = 0.271).

There was a significant non-linear correlation between T concentrations in adult plasma and egg yolks (SI appendix, Fig. S2A: LM: F_1,14_ = 5.263, p = 0.038), where concentrations of yolk T were lowest in eggs originating from females with intermediate levels of plasma T, but did not correlate with clutch size (SI appendix, Fig. S2B: F_1,14_ = 0.032, p = 0.862). In contrast, adult female plasma E_2_ concentrations were not correlated with E_2_ in the egg yolk (SI appendix, Fig. S2C: LM: F_1,21_ = 0.908, p = 0.351), but as clutch size increased, yolk E_2_ concentrations significantly decreased (SI Appendix, Fig. S2D: LM: F_1,21_ = 4.945, p = 0.037). The maternal E_2_:T ratio showed a non-linear relationship with the E_2_:T ratio in the egg yolk (SI Appendix, Fig. S2E: F_1,14_ = 6.493, p = 0.023), and was not correlated with clutch size (SI Appendix, Fig. S2F: F_1,14_ = 1.682, p = 0.215).

Immediately after oviposition, the clutches of these 26 females and two others (n = 28) were relocated into an *in-situ* experimental hatchery that was protected from terrestrial predation, yet exposed to natural sand and weather conditions. We buried clutches at a depth of 55 cm to standardise the thermal incubation environment. We confirmed the standardised thermal environment using data loggers placed at the centre of the clutch (mean thermosensitive period temperature = 30.02 ± 0.05 (SE) °C, SI Appendix, Fig. S3). The small amount of temperature variation observed was explained by differences in clutch size (F_1,26_ = 4.418, p = 0.045), resulting from increased metabolic heat produced from more developing embryos in larger clutches^34^. Assuming the pivotal temperature of this population to be 29 °C, as has previously been used for this population^4^, this incubation temperature would produce 12.89 ± 0.01 (SE) % male offspring if temperature was the sole determinant of sex ratio (Fig. 1A). Incubation duration, the time between oviposition and neonate emergence, is also often used as a proxy to predict offspring sex ratios (e.g. r^2^ = 0.73 in nests in Brazil^35^) and was recorded for each clutch^35^. Using the established logistic relationship between incubation duration and sex ratio observed in loggerhead turtles from Kyparissia, Greece (Fig. 1B, the closest location where the relationship between incubation duration and offspring sex ratio has been quantified for loggerhead turtles), the predicted sex ratio of our study clutches would be 47.5 ± 6 (SE) % males^36^. This suggests that levels of sex ratio variation are far greater than those we would expect from temperature alone.

**Fig. 1:**
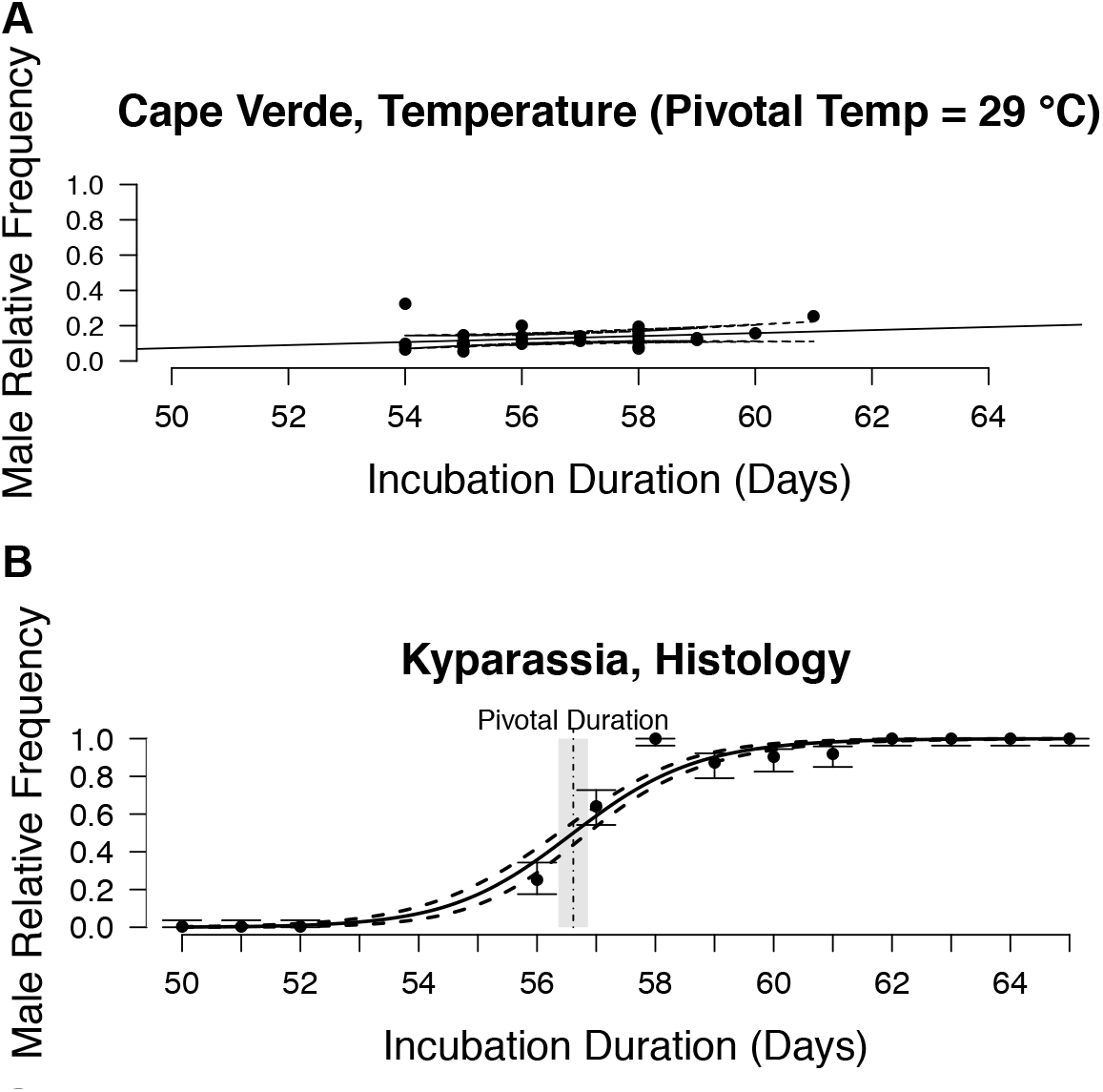
Sex ratios of study clutches A) as would be expected with a pivotal temperature of 29 °C B) based on the relationship between incubation duration and clutch sex ratios in Kyparassia (Greece)^36^

While the incubation duration represents a reasonable proxy for estimating the sex ratio of sea turtle offspring, currently the only accurate method to resolve individual sex requires sacrificing neonates and histological examination - a limiting factor for endangered populations^35,37^. However, we developed a new method to ascertain individual sex without the need to sacrifice animals. After taking 100 – 150 μl of blood from 365 offspring from 28 clutches after emergence (mean offspring per clutch = 13 ± 4 (SE)), we measured plasma hormone concentrations using ELISA. Hatchling hormone levels varied among individuals (SI Appendix, Table S1) and among clutches, with the average E_2_:T ratio of clutches ranging from 1.06 ± 0.13 (SE) to 3.56 ± 0.68 (SE). We used affinity propagation clustering (APC) on hatchling E_2_:T ratios guided by incubation duration to identify clusters of individuals with a similar hormonal phenotype. APC iteratively considers the similarity of a data point to its neighbours. Importantly, it does not require the number of possible clusters to be defined *a priori,* as is necessary for other clustering approaches such as k-means^38^. We identified three APC clusters (Fig. 2A). Two of these originate from clutches with short incubation durations, the classic trait of female neonates, and were distinguished by differences in their mean E_2_:T ratio (SI Appendix, Fig. S4, Cluster 1: mean = 4.45 ± 0.26 (SE), Cluster 2: mean = 1.72 ± 0.05 (SE), t-test: df = 44.08, t = 10.273, p < 0.001). The third group is formed by individuals from clutches with longer incubation durations (t-test: df = 299.3, t = −32.933, p < 0.001) and a low E_2_:T ratio (SI Appendix, Fig. S4, mean = 1.52 ± 0.06 (SE)), the characteristics of male sea turtle neonates.

**Fig. 2:**
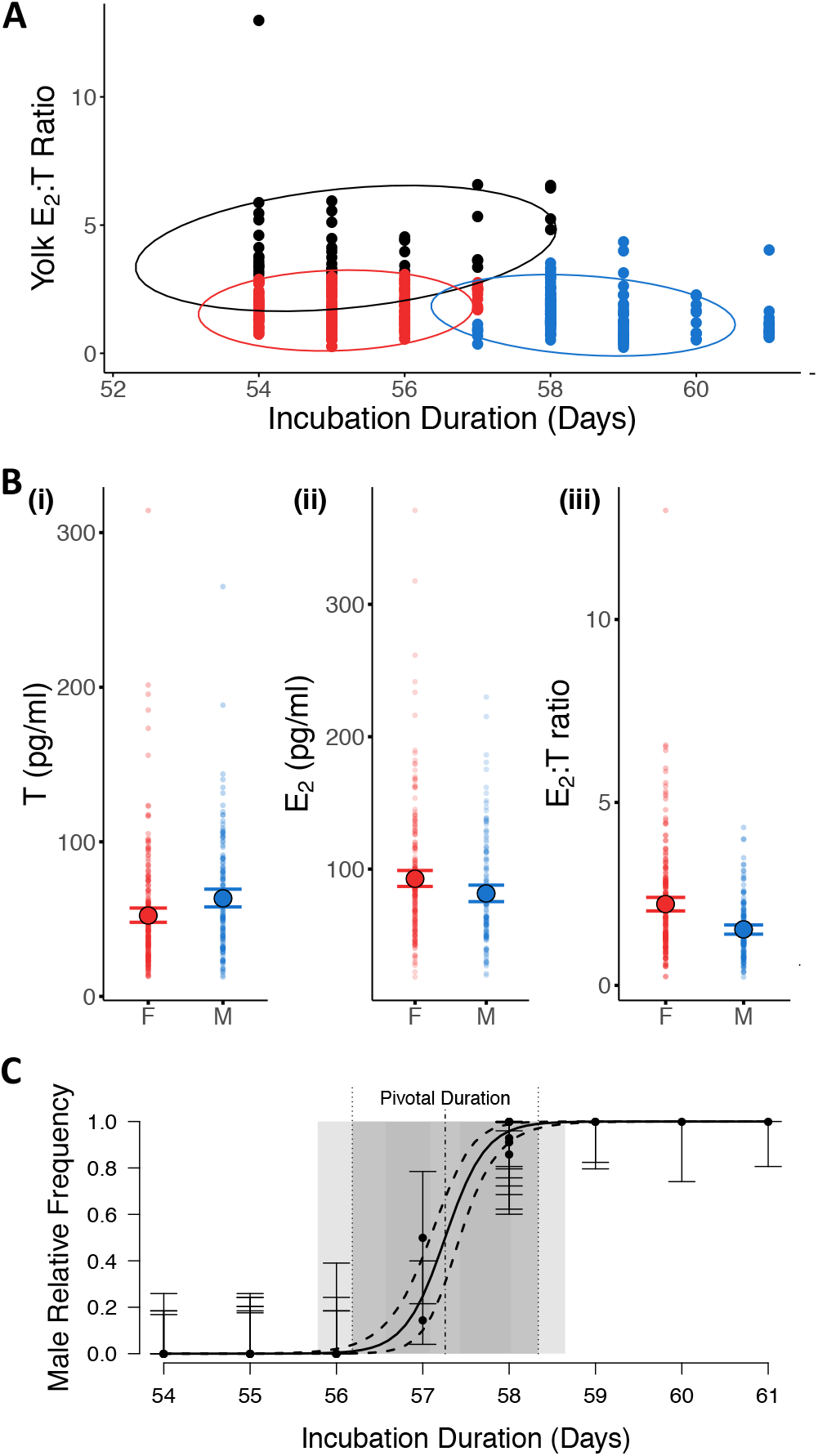
Individual sex as calculated by affinity propagation clustering (APC). A) APC identifies three different clusters based on individual E_2_:T ratio and clutch incubation duration. These clusters equate to female (red and black) and male (blue) offspring; B) Significant differences in the concentrations of T (F_1,60_ = 10.673, p = 0.002), E_2_ (F_1,57_ = 7.521, p = 0.008) and the E_2_:T ratio (F_1,48_ = 28.652, p < 0.001) between male and female offspring (mean, 95% confidence intervals and raw data are shown); C) Frequency of male offspring estimated by APC in relation to incubation duration. The pivotal duration was estimated at 57.25 days.

Several positive theoretical controls were used to confirm this method since neonate sacrificing is not possible. First, linear mixed effect models (LMM) using clutch ID as a random factor revealed significant differences in hormone levels between the two sexes, that were directly comparable to previous studies in which individuals’ sex was confirmed through histology^39,40^. As expected, T levels were higher in males (Fig. 2Bi, LMM: F_1,60_ = 10.673, p = 0.002, mean = 63.63 ± 2.89 (SE) pg/ml) than in females (mean = 52.54 ± 2.34 (SE) pg/ml), and conversely E_2_ levels were higher in females (Fig. 2Bii, LMM: F_1,57_ = 7.521, p = 0.008, mean = 92.94 ± 3.06 (SE) pg/ml) than in males (mean = 81.66 ± 3.16 (SE) pg/ml), as was the overall E_2_:T ratio (Fig. 2Biii, LMM: F_1,48_ = 28.652, p < 0.001, females: mean = 2.22 ± 0.09 (SE); males: mean = 1.52 ± 0.06 (SE)). LMMs did not detect any difference in weight (F_1, 348_ = 0.024, p = 0.878) or size (F_1, 218_ = 0.766, p = 0.382) between the sexes, as would be expected under these conditions by the Charnov-Bull theory^14^. Second, by combining individual offspring sex into an estimate of clutch sex ratio, and comparing this to the incubation duration, we found the specific logistic regression curve that characterises incubation durations in Type Ia TSD species (Fig. 2C). The pivotal duration was fitted to a value of 57.25 days (95% CIs: 57.09, 57.43), with a transitional range of incubation durations of 2.15 days (95% CIs: 1.52, 2.77). Importantly, if individual sex were incorrectly assigned, this distinctive logistic regression curve of TSD species would not be seen. With this method, we determined that clutch sex ratios were on average 40.49 ± 8.98 (SE) % male. This suggests 26.1% more males and far more variation in clutch sex ratio than would be expected based on incubation temperatures alone. Our sex ratio estimate is slightly below (7.1%) that estimated from parameters based on incubation durations in Kyparissia, suggesting population differences in development rate exist, likely as a result of different average pivotal temperatures among rookeries.

After establishing that the inter-clutch variation in sex ratio (and also in incubation duration, see SI Appendix Supplementary Analysis) was too great to be produced by temperature alone, we tested whether metabolic heat and/or maternal hormone transfer in the yolk predicted incubation duration and the estimated sex ratio. Yolk T correlated negatively with both incubation duration (LM, F_1,22_ = 10.624, p = 0.003) and the proportion of males produced within a clutch (Fig. 3A, Binomial generalised linear mixed effect models (GLMM), x^2^ = 4.371, df = 1, p = 0.037), but metabolic heat had no detectable effect (incubation duration model: F_1,22_ = 2.436, p = 0.133, sex ratio model: x^2^ = 2.111, df = 1, p = 0.146). There was no relationship between yolk E_2_ and incubation duration or clutch sex ratio (Fig. 3B, incubation duration: F_1,23_ = 3.169, p = 0.088, sex ratio: x^2^ = 0.183, df = 1, p = 0.669), yet the yolk E_2_:T ratio showed a non-linear relationship with both incubation duration (Fig. 3C, F_1,21_ = 12.882, p = 0.002) and sex ratio independently of temperature (x^2^ = 39.319, df = 2, p < 0.001). A maximum incubation duration of 57.2 days was observed at an equal hormone ratio (E_2_:T of 1.05, y = −7.8x^2^ + 16.3x + 48.7) with the highest levels of male offspring developing at this point. Because of the presence of a possible outlier with a high yolk E_2_:T ratio, we re-analysed the data without this point. The same patterns remained with significant non-linear relationships between yolk E_2_:T ratios and incubation duration (F_1,20_ = 6.292, p = 0.021) as well as between yolk E_2_:T ratios and hatchling sex ratios (x^2^ = 7.584, df = 2, p = 0.022).

**Fig. 3:**
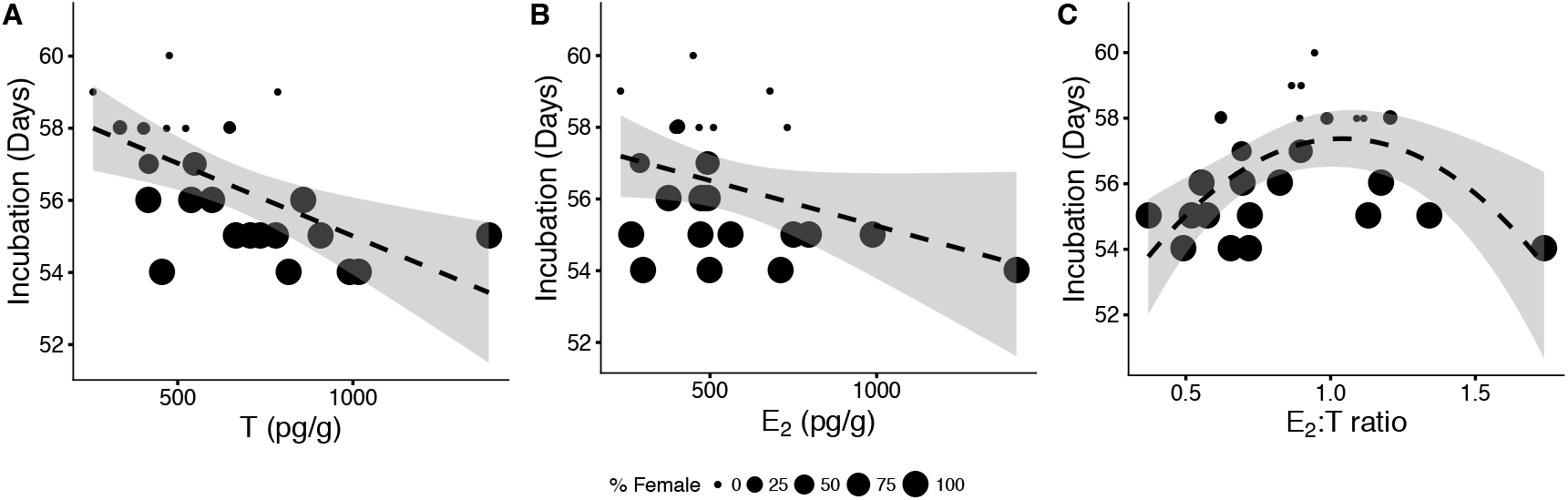
Relationship between maternally derived A) Testosterone T (F_1,22_ = 10.624, p = 0.003), B) Estradiol E_2_ (F_1,23_ = 3.169, p = 0.088) and C) E_2_:T ratio (F_1,21_ = 12.882, p = 0.002) and concentrations within egg yolks and incubation duration. Size of data points relates to the sex ratio as proportion of females as determined by APC.

Male offspring production was highest when maternal investment of E_2_ and T to the yolk was equal. Asking whether the production of either sex is more costly in terms of total hormone investment, we compared the total hormone concentration (E_2_ + T) with the overall E_2_:T ratio. This relationship was again non-linear, with total hormone investment being highest when the E_2_:T ratio was unequal (SI Appendix, Fig. S5A, LM: F_2,22_ = 4.951, p = 0.017), suggesting that producing females requires more maternal investment than males. The total hormone investment also showed a non-linear relationship with clutch size (SI Appendix, Fig S5B, log(E_2_ + T): F_2,22_ = 4.306, p = 0.026), with an initial increase in investment across clutch sizes between 65 and 75 eggs, after which investment declined with increasing clutch size.

Finally, to illustrate how maternal hormone transfer could impact population dynamics, we re-parameterised a previously published mathematical projection of neonate sex ratios for the Cape Verde population^4^, which assumed a fixed pivotal temperature of 29 °C. We made the simple assumption that the effect of maternally derived hormones on sex ratio is constant across a thermal gradient and applied the 26.1% observed difference in male offspring production for the coming century (Fig. 4). With a mechanism of this possible strength, the population is unlikely to reach the levels of extreme feminisation previously forecasted – instead of female production reaching over 97% in 2100, it is likely to instead reach approximately 71%. As it remains to be determined how maternal hormone transfer interacts with different incubation temperatures, this model only illustrates the potential importance of trans-generational hormone transfer for population dynamics.

**Fig. 4:**
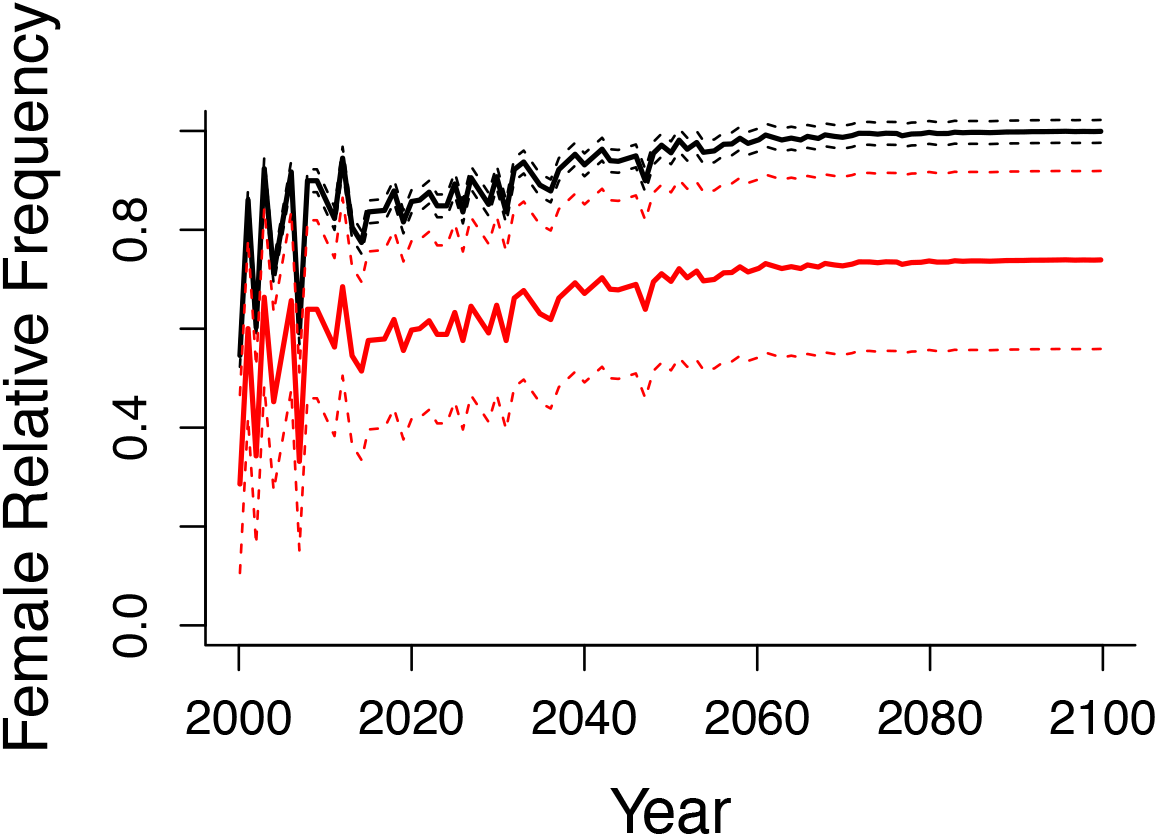
The population sex ratio of Cape Verde over the next century if it was determined by temperature alone ^4^ (black) and incorporating the effect of hormones observed here on the sex determining mechanism (red) along with the 95% confidence intervals.

## Discussion

It is widely speculated that global warming can drive species with temperature-dependent sex determination to extinction because of the over-production of one sex. But, given the many considerable historical shifts in climate experienced by TSD species, they are likely to have evolved behavioural and/or physiological mechanisms to avoid unviable biases in offspring sex ratio^5^. By experimentally standardising the thermal environment of loggerhead sea turtle nests *in-situ*, we investigated whether maternally derived hormones correlate with offspring sex independently of temperature. First, we developed a non-lethal method to determine the sex of neonates upon their nest emergence, using affinity propagation clustering based on individual circulatory sex steroid hormones and their incubation duration. With this method, we found a non-linear relationship between the clutch sex ratios and the ratios of maternally derived E_2_:T within the egg yolks under standardised thermal conditions. Low concentrations of equal investment in both hormones within the yolk maximise the production of male offspring, while increasing the concentration of either E_2_ or T, along with overall hormone investment, feminises the clutches. Re-parameterising an existing model that predicts sex ratio biases in response to climate change demonstrated that this trans-generational mechanism could prevent the predicted extreme feminisation of loggerhead turtles in Cabo Verde.

To date, an inability to determine neonate sex non-lethally has constrained the study of TSD mechanisms in endangered sea turtles (but see^32^). A clustering approach that identifies individuals with similar phenotypes (here hormone profiles) that match control traits of male and female offspring (incubation duration) overcame this problem. Using E_2_:T thresholds to define neonate sex has been verified with histological analysis in loggerhead^40^ and green^39^ turtles, but as E_2_:T levels vary considerably among clutches, it is difficult to delineate a population level threshold *a priori*. Using an APC method guided by incubation duration to group hormone profiles, a common proxy for sex ratio, we avoid the need to define thresholds and, importantly, the need to sacrifice individuals^35^. Because ethically we cannot validate this method further, we rely on strong indirect evidence, such as (i) the identified significant difference in circulating E_2_:T ratios of male and female offspring (ii) the pivotal duration both match those reported in other studies^36,39–42^, and (iii) the relationship between sex ratio and incubation duration fits the known logistic regression curve observed in Type Ia TSD species. We anticipate that this non-lethal approach will prove an invaluable tool for both research on TSD in sea turtle species and also for wider conservation.

Despite the standardised thermal environment of clutches within this experiment, high levels of variation in incubation duration and sex ratio were observed among nests, both of which correlated with maternally derived hormones within the egg yolk. The relationship between the yolk E_2_:T ratio and clutch sex ratio was best described by a quadratic curve, centred on an equal concentration of both hormones and ranging from 0.37 to 1.73. When maternal investment of E_2_ and T was equal, incubation durations were long, and males were produced. If hormone investment was biased in either direction, sex ratios became increasingly feminised. The effects of elevated levels of both E_2_ and T on sex ratios in this study are consistent with experimental manipulation of these hormones in other species^2,20,22^. In addition, the non-linear relationship explains studies where exogenous application of E_2_ has unexpectedly produced male hatchlings (e.g.^43, 44^). In such cases, E_2_ application, combined with existing maternal contributions, may have resulted in shifting the E_2_:T ratio within the eggs closer to one, forcing male development. Interestingly, in all other reptiles for which data are available, the E_2_:T ratios that are transferred to the yolk consistently remain below or above the ratio of one^45^. Thus, our study shows how maternal transfer of both hormones can influence the feminisation process of reptiles under natural conditions.

Total hormone concentrations within the yolk were lowest at an equal, male producing, E_2_:T ratio. If this ratio departed from 1:1 in either direction, total concentrations of yolk hormones increased. As E_2_ and T positively correlate within the egg, if investment in either hormone is elevated, there is an associated increase in the other. The outcome is that greater investment is required to skew E_2_:T ratios in a manner that favours the production of female offspring. This maternal investment provides initial hormonal substrate with which to prime the reactions required to activate molecular pathways that result in female gonad development. When E_2_:T ratios are skewed, and total hormone concentrations are high, feminisation is easily achieved through either the presence of E_2_ directly, or by the synthesis of E_2_ from its precursor, T, by the aromatase enzyme. When E_2_ and T are in equilibrium, low concentrations of E_2_ are not sufficient to feminise the clutch. However, product-feedback inhibition of aromatase receptors likely prevents further E_2_ being synthesised from T, and consequently male offspring are produced.

There is no doubt that temperature is the primary determinant of sex in TSD species, but studies have repeatedly recorded variation in sex ratios among clutches exposed to similar temperature regimes^32,46^. Maternally-derived E_2_ and T can affect the same developmental pathways as temperature, and explain some of this variation by priming reactions required to initiate female producing pathways^2,22^. Interestingly, the effects of these hormones on sex determination vary by experiment and species, likely as the results of adaptation to local nesting conditions (e.g.^43, 44^). However, here we show that in sea turtles, a clutch specific threshold exist for feminisation that is the product of an interaction between temperature and maternal hormone transfer. A shift towards an equal E_2_:T ratio and lower maternal investment will increase the pivotal temperature away from the feminisation threshold, and consequently warmer temperatures would be required to feminise a clutch. This aligns with the sex ratios observed within this study, which contained 26.1% more males than expected from a pivotal temperature of 29 °C. However, this mechanism will be constrained by physiological limits of maternal hormone investment. Our findings provide mechanistic explanations for the high inter-clutch variation that has been observed in TSD systems (eg^11, 46^), and also clarify occasions where female offspring have been produced at assumed male producing temperatures, or vice versa^32^.

Two maternal traits show a relationship with levels of hormone transfer to the clutch. Firstly, T concentrations within the yolk correlated non-linearly with those in maternal plasma. Disentangling the cause of such a relationship is complex as it is likely to result from multiple physiological cascades^47^. As vitellogenesis and follicular development in sea turtles occurs prior to migration, it is also likely that T concentrations in maternal plasma varies when yolks is formed^48^. However, this relationship does allow us to link T investment to maternal state. Should maternal T vary in response to environmental cues, as in the spined toad (*Bufo spinosus*), it may allow nesting females to plastically match the individual development threshold of feminisation to the ambient temperature, and maintain more constant sex ratios across a nesting season^49^. Similar differences in the maternally-transferred E_2_:T ratio in the egg yolk of a population of painted turtles resulted in a seasonal shift of the pivotal temperature, albeit in a direction that accelerated female production as temperatures increased^16^. Secondly, total E_2_ investment within eggs decreased as clutch size increased, and total hormone concentrations were low in large clutches. Thus, in large clutches with more metabolic heat production, the developmental threshold of feminisation is increased – minimising sex bias. We infer from these results that there are two distinct mechanisms that can affect the ratio of E_2_:T within the yolk, which explains how elevated investment in either hormone can lead to feminisation. There is considerable variation in circulating T and E_2_ levels between sea turtle populations and species (SI Appendix, Table S1), which may suggest an element of local adaptation in response to environmental conditions, and a heritable component of baseline physiological levels^50^.

TSD species will require behavioural and/or physiological responses to maintain viable sex ratios in the face of future climate change. Here, we highlight a previously under-considered physiological mechanism for individual variation in the TSD process within sea turtle species. There is a need for management plans that use temperature-based models to predict future sex ratios to account for maternal hormonal influence, as this will have considerable implications for population dynamics.

## Methods

### Sample Collection

We studied nesting loggerhead sea turtles on the island of Boavista, part of the Cabo Verde archipelago in the eastern Atlantic. The sampling site (15**°**58’18.6”N, 22**°**48’06.2”W) is a 400 m stretch of coastline on the southern tip of this island. Twenty-eight nesting females were sampled between 17 July and 1 August 2017. Immediately after oviposition, females were individually marked with PIT (AVID) and metal (Inconel) tags^51^. Blood samples of 1-4 ml in volume were collected from the dorsal cervical sinus of 26 females using a 40 mm, 21-gauge needle and 5 ml syringe, and stored within lithium heparin containers. Finally, curved carapace length (CCL) and width (CCW) were measured (± 0.1 cm).

The clutches of these turtles (containing 83 ± 3 (SE) eggs) were relocated to an experimental hatchery protected from predation, situated on the nesting beach. At this point, up to two eggs from the 28 clutches were removed from each clutch for yolk hormone analysis, and the rest of the clutch was buried at a depth of 55 cm. By using a standard depth, temperature was controlled for, while maintaining an otherwise natural environment. A TinyTag™ temperature logger was placed at the centre of each clutch, programmed to take a reading every 15 minutes throughout the incubation period (accuracy ± 0.2 °C). As anticipated, the uniform depth standardised the incubation temperature of the nests to 30.05 ± 0.05 (SE) °C during the middle third of incubation, the period where embryo sex is established. This variation in temperature is extremely conserved, and is representative of the thermal variation produced within treatments under controlled laboratory incubations^52, 53^.

Upon emergence, twenty hatchlings were randomly selected for blood sampling (100 – 150 μl) from the dorsal cervical sinus, using a 26-gauge needle and 1 ml syringe^54^. Samples were stored within lithium heparin coated tubes. Notch-to-notch straight carapace length (SCL) and, width (SCW) were measured using digital callipers (± 0.01 mm), and weight was measured with a digital scale (± 0.1 g)).

The blood samples of both the adults and offspring were refrigerated for up to 48 h before being centrifuged to extract plasma. Egg yolks were separated from the albumen, and all samples were stored at −20 °C until extraction.

### Hormone extraction

Commercially available Enzyme-Linked Immunosorbent Assay (ELISA) kits for both E_2_ (Catalogue # ADI-900-174, ENZO Life Sciences) and T (Catalogue # ADI-900-065) were used to measure steroid levels in all samples. Details for hormone extraction protocols are given in SI methods. Not all blood samples had sufficient volume for hormone extraction. Consequently, we extracted E_2_ from 24 adults and 388 hatchling blood samples, and T from 19 adult and 367 hatchling blood samples. This provided us with E_2_:T ratios for 18 adult females, and 365 hatchlings. E_2_, and T were successfully extracted from the yolks of 26 out of the 28 sampled clutches. One yolk T measurement was removed as an outlier, being more than three standard deviations from the mean (mean: 741.26 ± 502.29 (SD) pg/g, outlier: 2882.05 pg/g).

### Statistical Analyses

All analyses were conducted with R 3.3.3, using the R packages *lme4* and *lmerTest* for fitting linear mixed models (LMMs) and generalized linear mixed models (GLMMs). A paired t-test was used to compare intra-clutch E_2_ and T levels between two eggs in a subset of clutches (n = 13), to test whether that there was variation in hormone investment between eggs in a clutch. As there was no difference between eggs from the same clutch, for subsequent analyses the average hormone was used where possible, while a single egg was used for the remainder of the clutches. Correlations between E_2_ and T in female plasma and yolks, the effect of clutch size on temperature, and the effects of metabolic heat and maternally derived hormones on incubation duration were tested using general linear models (LM). A non-linear relationship between the E_2_:T ratio on incubation duration was fitted using a quadratic curve. Similarly, when considering the relationship between E_2_:T and total hormone investment, we also fit a quadratic model. LMs were also used to estimate the correlation of clutch size and plasma hormone concentrations with yolk hormone concentrations.

We used Algorithm Propagation Clustering (APC) to identify individual sex, using the R package *apcluster*^38^. Cluster assignment was made based on the plasma E_2_:T ratio of hatchlings, guided by their incubation duration. Determining neonate sea turtle sex using the E_2_:T ratio has previously been extremely accurate (96% and 96.7% respectively) for artificially incubated eggs of loggerhead and green sea turtles^39,40^ that were ultimately sacrificed for verification. Since variation likely exists among rookeries, those thresholds however cannot be blindly applied to new populations. T-tests were used to compare hormone levels of putative male and female hatchlings. A response curve of these estimated sex ratios to incubation duration was produced using the logistic equation function of the R package *embryogrowth* to further verify the accuracy of our non-lethal sexing method.

After identifying the sex of individuals, LMMs were used to compare individual size and weight between the sexes and the APC clusters. Finally, we used binomial GLMMs to determine whether individual hatchling sex was predicted by maternal hormone investment or temperature. For all LMM and GLMM analyses, clutch was included as a random factor to account for individual variation. Model selection was based on AIC criteria, using a likelihood ratio tests to select for the best models. P-values of the selected models were obtained by with the *car* R package, and models were verified for over-dispersion.

Thermal estimates of sex ratio were calculated using the equation first presented by Girondot in 1999 with the R package *embryogrowth*, under an assumed pivotal temperature of 29 °C^55^. Estimates of sex ratio based on incubation duration were made based on data from a study on a neighbouring loggerhead sea turtle population that nests in Kyparissia, Greece, and was confirmed with histology^36^. To generate an illustrative model that compared the results of our study with future predictions based on temperature alone, we extracted data from a previously published study predicting sex ratios until 2100 based on a fixed pivotal temperature of 29 °C alone. We then compared our observed mean clutch sex ratio to that expected from a pivotal temperature of 29 °C, and added the difference, along with 95% confidence intervals, to the original prediction.

## Acknowledgements

This work was carried out as part of The Turtle Project, a citizen science project dedicated to sea turtle research and conservation. We thank volunteers and staff from Turtle Foundation in particular for facilitating this research. Particular thanks go to S. Cameron for considerable help during data collection. E.L was supported by a studentship from the London Natural Environment Research Council Doctoral Training Partnership (grant NE/L002485/1). This study was supported by a National Geographic grant (NGS-59158R-19), Natural Environment Research Council (NE/V001469/1) as well as Queen Mary University of London Funds allocated to CE. We also acknowledge the support from “The Future Ocean’ Excellence Initiative by the Deutsche Forschungsgemeinschaft (DFG) that granted a ‘Capacity building grant’ to C.E. The authors would like to thank the Eizaguirre lab as well as Dr. José María Martín-Durán and Dr. Gail Schoefield for their feedback on previous versions of the manuscript.

## Author Contributions

E.L. and C.E. designed the experiment. T.R. facilitated the fieldwork. E.L. and C.E. conducted the fieldwork. E.L. analysed the data and drafted the initial manuscript, with feedback from C.E.. All authors approved the manuscript.

