## Supplementary Material for "Maternally derived sex steroid hormones impact sex ratios of loggerhead sea turtles"

**This PDF file includes:**

Supplementary Material and Methods

Supplementary Analysis

Figure S1

Figure S2

Figure S3

Figure S4

Figure S5

Table S1

References for SI reference citations

Supplementary Materials and Methods

**1. Hormone Extraction Protocol**

Both testosterone (T) and oestradiol (E_2_) were extracted from adult and hatchling plasma samples following commercial Enzyme-Linked Immunosorbent Assay (ELISA) kit protocols (E_2_: Catalogue # ADI-900-174, T: Catalogue # ADI-900-065, ENZO Life Sciences). Anhydrous diethyl ether was added at a ratio of 5:1 to a 40-200 μl plasma sample. After homogenizing and snap freezing, the diethyl ether fraction was decanted into a fresh test tube containing either 0.5 or 1 ml of distilled water and homogenised once more. The resulting organic phase was removed and evaporated for two hours using a speed vacuum. Samples were then rehydrated using 250 μl of appropriate assay buffer from the ELISA kits, and frozen until assayed. Yolk hormones were extracted following the protocol from Schwabl^1^ with the exception of the final hexane phase, which was not conducted. After extraction, samples were reconstituted in 250 μl of appropriate assay buffer.

Extraction efficiencies were determined for both hormones by dividing either adult plasma samples (E_2_: n = 6, T: n = 5) or yolk samples (E_2_: n = 6, T: n = 5) into two aliquots. One of these was spiked with a known concentration of hormone (E_2_: 272 pg/ml, T: 400 pg/ml,) prior to extraction. Efficiency was determined by calculating the difference between the spiked and non-spiked samples compared to the known spike quantity (Plasma: E_2_: 54.5 ± 10.5 (SE) %, T: 43.9 ± 2.8 (SE) %, Yolk: E_2_: 77.9 ± 6 (SE)%, T: 120.2 ± 9.8 (SE) %).

Serially diluted standards of known hormone concentration were prepared (E_2_ = 7, T = 5) according to the kit’s protocol, producing standard curves ranging from 1000 – 15.6 pg/ml (E_2_) and 2000 – 7.81 pg/ml (T). Samples were run in duplicate, and hormone concentrations were calculated using a curve-fitting program (MARS). All yolk samples were run on a single plate, with average intra-assay coefficients of variation (CV) being 10.63 ± 1.24 (SE) % for E_2_ and 9.89 ± 1.32 (SE) % for T. Hatchling plasma samples were run across six plates for each hormone. Intra-assay CVs were 13.5 ± 1.3 (SE) % for E_2_ and 17.05 ± 2.9 (SE)% for T.

**Supplementary Analysis:**

**1. Circulating E_2_ and T within adult plasma**

To support our observation that the levels of circulating E_2_ and T within adult plasma are significantly correlated, we augmented our dataset by also including samples collected from nesting turtles in the same location in 2016 (2016: n = 75, total including this dataset: n = 92). With this larger sample size, we confirmed the relationship observed within this study, showing a strong positive correlation between the two hormones (F_1,91_ = 93.527, p < 0.001).

**2. Relationship between metabolic heat and incubation duration**

There is a well-known link between heat and developmental rates^2^. Thus we quantified how metabolic heat influenced incubation duration within this study. Clutch size positively correlated with temperature as a result of metabolic heat production (Fig. S5A, F_1,26_ = 4.42, p = 0.05), and elevated metabolic heat increased embryonic development rates, and resulted in shorter incubation durations (Fig. S5B, F_1,26_ = 4.31, p = 0.05). Based on the slope estimate of this model, if metabolic heat was the sole determinant of variation in incubation duration within our study clutches, the range of temperatures recorded should cause the incubation duration to vary by a maximum of three days (95% CI = 2.4, mean: 56.63 ± 0.14 (SE), min = 55.34; max = 58.44 days). Yet, observed incubation durations ranged across seven days, from 54 to 61 days (mean: 56.57 ± 0.37 (SE)), implying that factors other than metabolic heat production associated with clutch size contributed to variation in rates of embryonic development – in this case, intrinsic egg characteristics.

| A  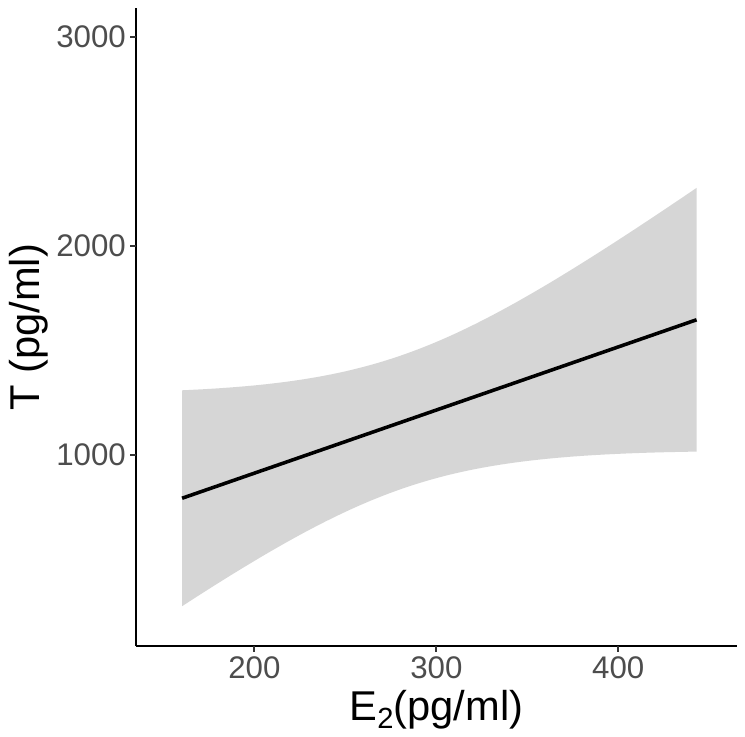 | B  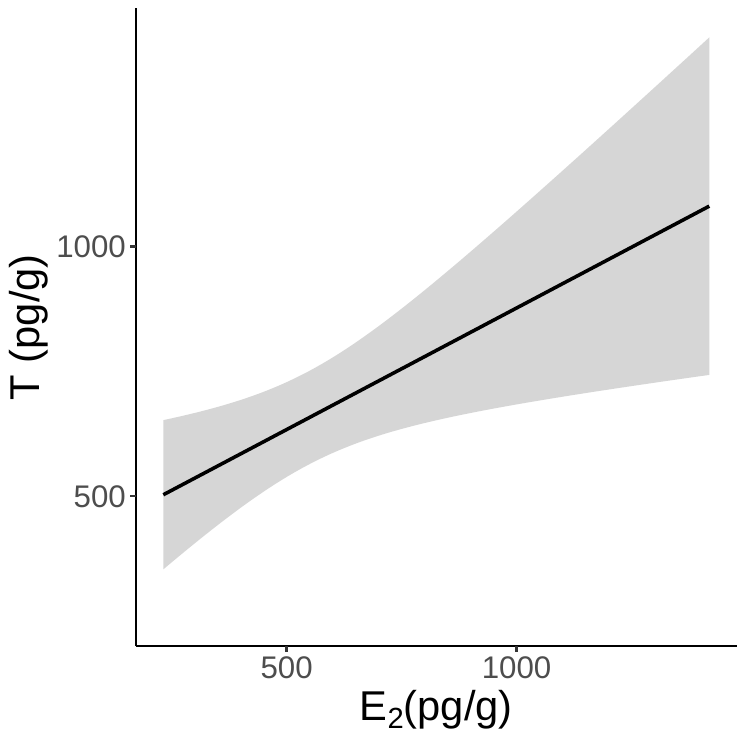 |
| --- | --- |

Fig. S1. Significant positive correlations between E_2_ and T in both A) adult plasma (F_1,16_ = 4.608, p = 0.048) and B) egg yolks (F_1,23_ = 7.338, p = 0.013)

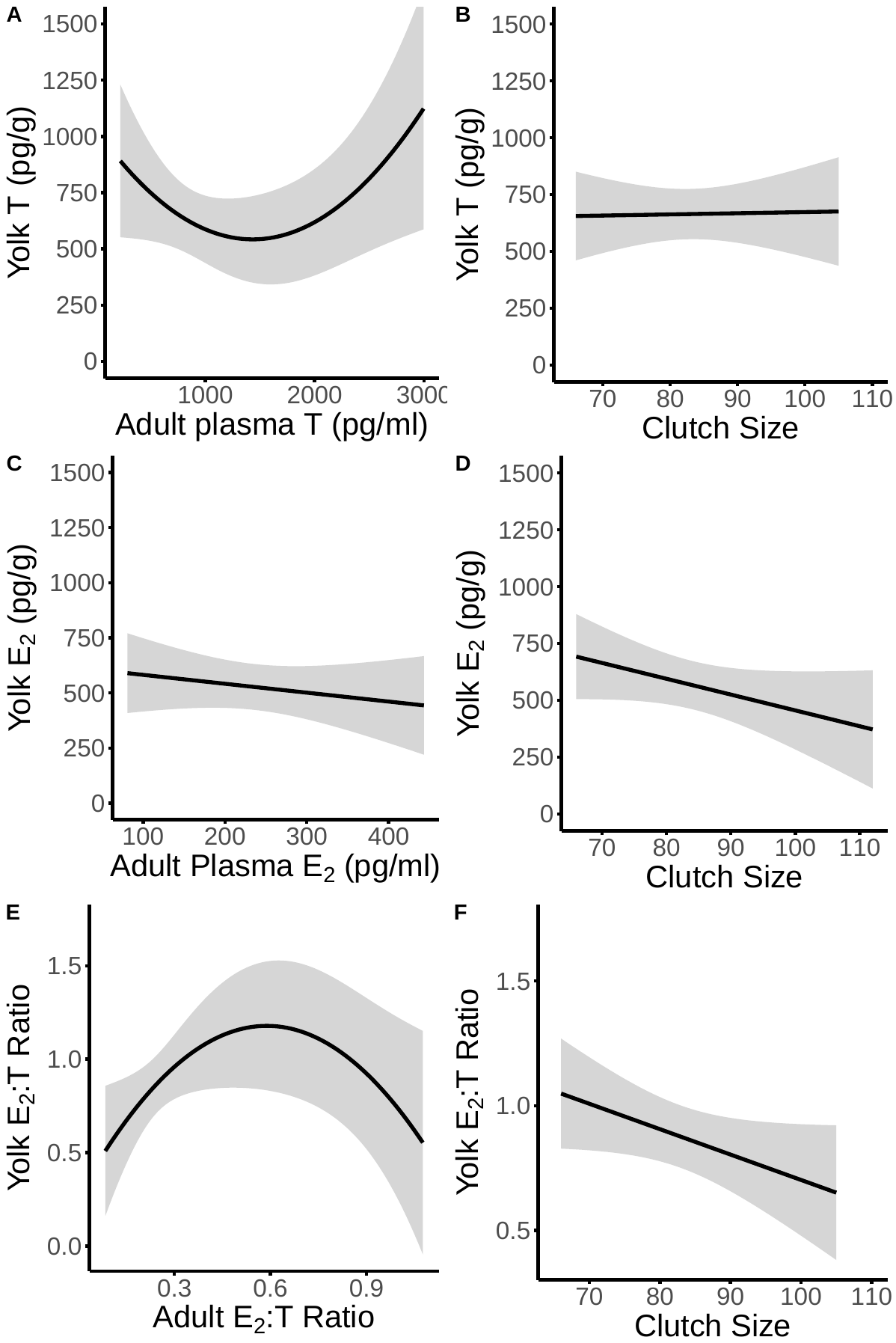

Fig. S2. The relationship between both adult circulating hormone concentrations and clutch size on yolk hormone concentrations. A) Yolk T has a significant, non-linear relationship with circulating adult T (F_1,14_ = 5.263, p = 0.038); B) Clutch size has no effect on yolk T concentrations (F_1,14_ = 0.032, p = 0.862); C) There is no relationship between Yolk E_2_ and the concentrations found in circulating adult plasma (F_1,21_ = 0.908, p = 0.351); D) As clutch size increases, concentrations of E_2_ within the yolk decrease (F_1,21_ = 4.945, p = 0.037); E) There is a non-linear relationship between the E_2_:T ratio found within the yolk, and that within adult female plasma (F_1,14_ = 6.493, p = 0.023); F) As clutch size increases, the yolk E_2­:_T ratio decreases (F_1,14_ = 1.682, p = 0.215).

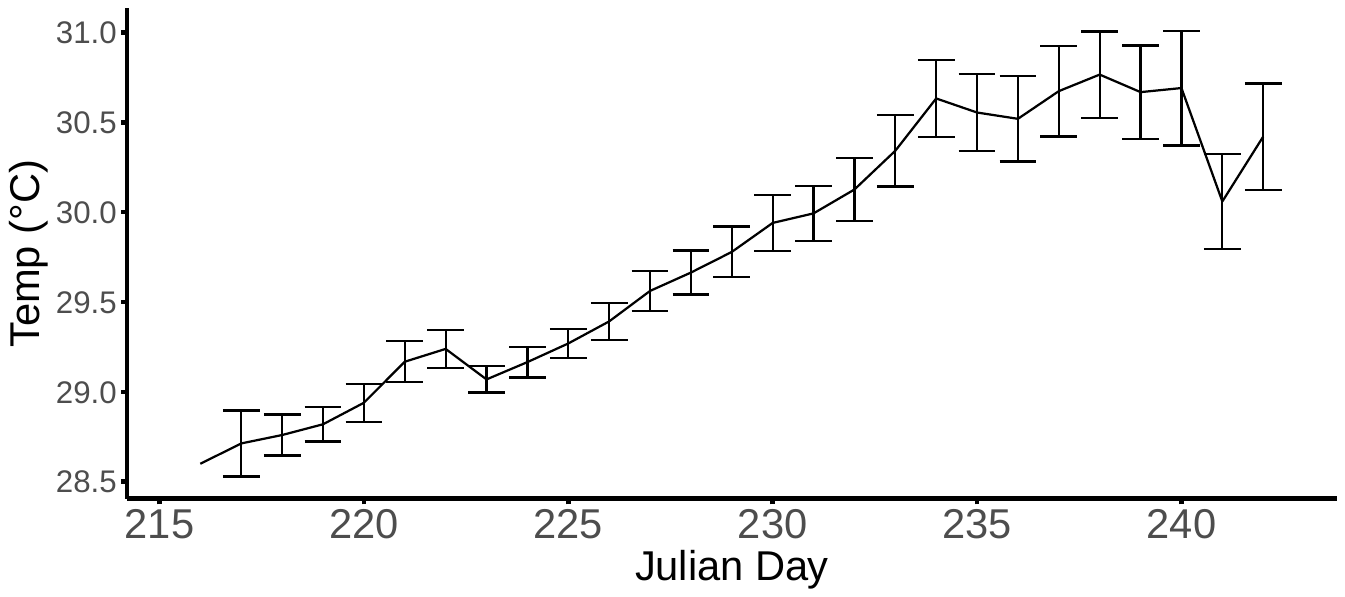

Fig. S3 Mean temperature ± 95% CI of 28 clutches of eggs through the thermosensitive period of incubation in 2017

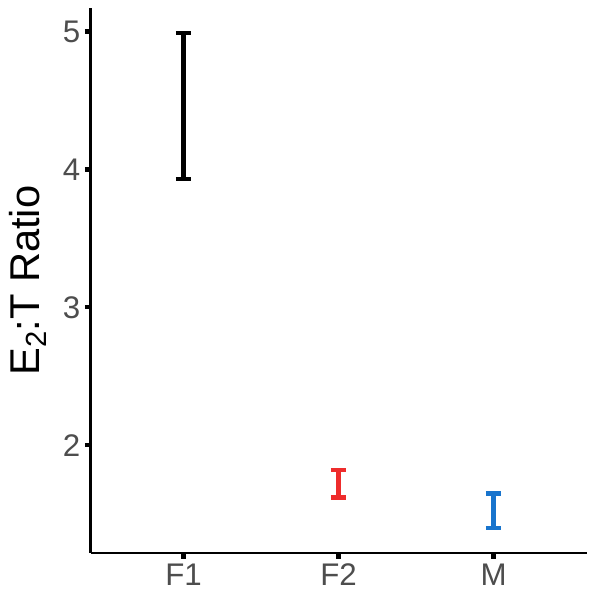

Figure S4: Differences in mean E_2_:T ratio (with 95% confidence intervals) between the three different clusters identified by APC clustering. Clusters F1 (n = 42) and F2 (n = 188) are assigned to be female from their E_2_:T ratio differences and short incubation duration. Cluster M (n = 151) is produced under the long incubation durations characteristic of male offspring, and have low E_2_:T ratios.

| A  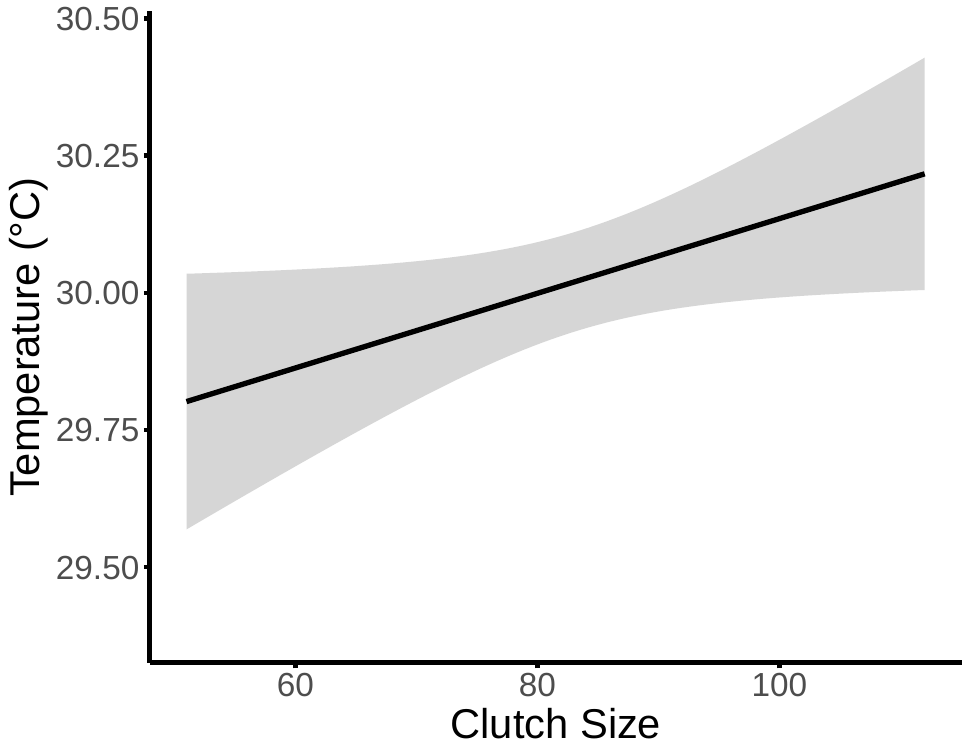 | B  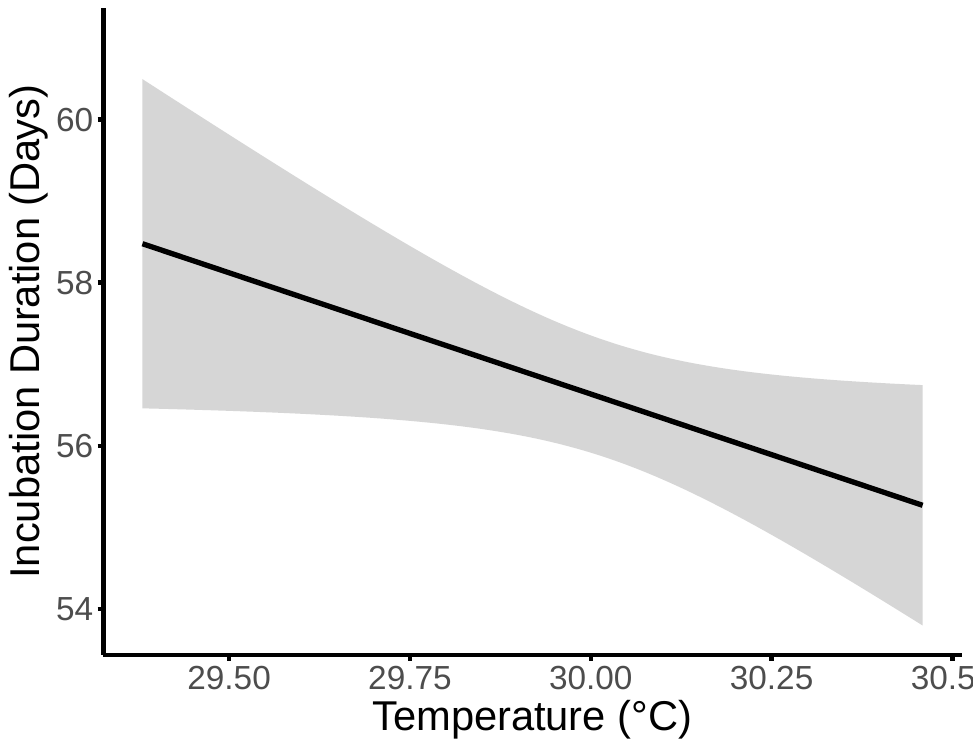 |
| --- | --- |

Fig. S5 Characteristic relationships between clutch size, temperature and rate of development were observed in study clutches. A) There was a significant positive relationship between clutch size and the temperature of the nest (F_1,26_ = 4.418, p = 0.045); B) Temperature had a negative influence on incubation duration (F_1,26_ = 4.311, p = 0.048)

Table S1. Adult and neonate plasma E_2_ and T concentrations and the E_2_:T ratio recorded in populations of sea turtles globally. Values from this study in bold. *Measurements taken from beginning of nesting season only. **Measurements taken from free-swimming, reproductively active female turtles at varying points within the nesting season.

| **Location** | **Species** | **n** | **Category** | **Sex** | **E_2_ pg/ml** | **T pg/ml** | **Ratio** |
| --- | --- | --- | --- | --- | --- | --- | --- |
| ***Adults*** |  |  |  |  |  |  |  |
| Oman^3^ | GT | 22 | nesting | F | undetected | 420 ± 40 (SE) | NA |
| Costa Rica^4^ | LB | 32 | nesting* | F | 190 ± 16.8 (SE) | 10180 ± 77000 (SE) | NA |
| Costa Rica^5^ | LB | 13 | nesting* | F | 53.30 ± 6.54 (SE) | 2224 ± 280 | NA |
| Australia^6^ | HB | 95 | nesting | F | 0 - 119 (range) | 0-7520 (range) | NA |
| Florida^7^ | LH | 38 | free-swimming** | F | 3.2 - 3723 (range) | 50 - 12900 (range) | NA |
| **Cape Verde** | **LH** | **26** | **nesting** | **F** | **235.8 ± 22.7 (SE)** | **1148.5 ± 148.6 (SE)** | **0.32 ± 0.05 (SE)** |
| ***Neonates*** |  |  |  |  |  |  |  |
| China^8^ | GT | 16 | post-emergence | M | 132 ± 37  (SD) | 186 ± 58  (SD) | 0.788 ± 0.338 (SD) |
| China^8^ | GT | 14 | post-emergence | F | 205 ± 50  (SD) | 105 ± 30  (SD) | 2 ± 0.438 (SD) |
| Japan^9^ | LH | 90 | post-emergence | NA | 0 - 50.2 (range) | 9.2 - 300.2 (range) | 0.01 - 1.24 (range) |
| North Carolina/  Florida^10^ | LH | 17 | post-emergence | M | 106 ± 15 (SD) | 215 ± 38 (SD) | 0.6 ± 0.1 (SD) |
| North Carolina/  Florida^10^ | LH | 11 | post-emergence | F | 198 ± 44 (SD) | 76 ± 13  (SD) | 2.7 ± 0.4 (SD) |
| **Cape Verde** | **LH** | **151** | **post-emergence** | **M** | **81.66 ± 3.16 (SE)** | **63.63 ± 2.9**  **(SE)** | **1.52 ± 0.06 (SE)** |
| **Cape Verde** | **LH** | **253** | **post-emergence** | **F** | **92.93 ± 3.06 (SE)** | **52.53 ± 2.34 (SE)** | **2.22 ± 0.09 (SE)** |
